# A viral infection reshapes Arabidopsis water management *via* root hydraulics, aquaporin downregulation and osmotic adjustment

**DOI:** 10.64898/2025.12.15.694488

**Authors:** Carlos Augusto Manacorda, Pablo Daniel Cáceres, Moira Romina Sutka, Gabriela Amodeo, Sebastián Asurmendi, Irene Baroli

## Abstract

The effect of plant viruses on root water relations and on how roots and shoots coordinate under infection remains poorly understood. Using a hydroponic *Arabidopsis thaliana*–Turnip mosaic virus (TuMV) pathosystem, we integrated biometric, anatomical, hydraulic, and gas-exchange measurements to dissect how viral infection reshapes root–shoot water relations. TuMV impaired root development, as reflected by an early plateau of primary root elongation. At the functional level, infected plants exhibited a decrease in root hydraulic conductance per unit root mass, concomitant with transcriptional downregulation of root aquaporin genes. Despite this, the relative contribution of aquaporin-mediated water transport, assessed *via* sodium azide inhibition, remained unchanged, indicating that the virus downregulates total hydraulic capacity without altering the apoplastic–symplastic partitioning of water flow. Gas-exchange analysis revealed a virus-induced decoupling between stomatal conductance and net CO₂ assimilation, resulting in a non-adaptive increase in intrinsic water-use efficiency. This loss of photosynthetic plasticity, combined with shoot-localized osmotic adjustment (more negative leaf osmotic potential and higher relative water content), points to a constrained, suboptimal physiological state. Multivariate analysis confirmed that variation in physiological traits largely drives phenotypic divergence between treatments. Together, these coordinated alterations, reduced root hydraulics, rigid gas-exchange relationships and passive hydraulic matching to a stunted shoot, depict plants locked into a low-performance equilibrium, poorly equipped to compete for water and carbon. This work reveals a systemic hydraulic–photosynthetic reconfiguration that could account for compromises in plant resilience and resource competitiveness.

**Highlight:** TuMV infection induces a coordinated whole-plant hydraulic reconfiguration characterized by premature growth arrest, reduced root hydraulic conductance, and decoupling of stomatal conductance from photosynthesis, resulting in a constrained physiological state.

## Introduction

Maintaining a proper water status when facing biotic and abiotic stressors is essential for plant growth and development and for ecosystem health (Torres-Ruiz et al., 2024). Plant viruses are an important source of worldwide agricultural losses in crops (Loebenstein, 2008; Scholthof et al., 2011), which threaten food security in developing countries and determine economic losses globally (Jones & Naidu, 2019). In this context and given that viral infections can alter leaf water relations (Aguilar & Lozano-Duran, 2022; Pollari et al., 2022), key open questions are whether, and how, the root hydraulic pathway in infected plants is coordinated with these demand-side changes. Addressing this requires a whole-plant hydraulic framework.

Plant hydraulics depends strongly on environmental conditions, especially on the water content of the surrounding soil and air, since water moves through the plant by a passive mechanism, involving diffusion and bulk flow, in the soil–plant–atmosphere continuum or SPAC (Passioura, 1982; Boyer, 1985). To dynamically adjust to soil water availability and evaporative demand in the short term, plants regulate stomatal opening, while in the long term they can modify the hydraulic properties of the root and shoot. Stomata represent the predominant resistance to water movement in the SPAC, with their aperture determining the rate of transpiration for a given evaporative demand (Buckley, 2019). In contrast, roots make up less than half of the plant hydraulic resistance and therefore roots have received less attention as sites of dynamic regulation of water transport. However, there is increasing evidence that plants can adjust root hydraulic resistance in response to environmental pressures (Maurel et al., 2010; Vadez, 2014). Furthermore, it has been shown that the hydraulic conductivity of roots correlates with the gas exchange capacity of the shoot, both under stress and under normal growth conditions (Sade et al., 2009; Vandeleur et al., 2014; Rodriguez-Dominguez & Brodribb, 2020).

The relative contribution of the cell layers of the root cortex to whole root hydraulics can be explained by the composite transport model proposed by Steudle (2000). To be absorbed from the soil and reach the root vasculature, water can pass through the tissues either through the cell walls, i. e., *via* the apoplastic pathway, which has a low resistance, or the cell-to-cell pathway, comprising plasmodesmata and biological membranes, with higher resistance. The magnitude of the osmotic and hydrostatic forces will determine which path is the primary contributor to radial water flow. The model allows for a coarse regulation of radial transport, involving the modulation of apoplastic barriers, such as xylem anatomy and the degree of endodermis suberization, and a fine regulation brought about by the abundance and activation state of aquaporins, which provide a major pathway for transmembrane water flow and can modulate root hydraulic conductance (Maurel et al., 2016). Although the lipid bilayer is permeable to water, the presence of aquaporins dramatically increases the permeability of the membrane that contains them (Chaumont & Tyerman, 2014). There is increasing evidence that the modification of aquaporin activity is an important trait in root water management. Changes in aquaporin gene expression, post-transcriptional modifications, protein-protein interactions and gating mechanisms impacts in plant water homeostasis, regulating dynamic changes of root, stem and leaf hydraulic conductivity (Vitali et al., 2015; Moshelion et al., 2015; Ozu et al., 2018).

Although the effects of abiotic stresses, especially of water deficit, on the hydraulic conductivity of the root, Lp_r_, have been extensively studied (Aroca et al., 2012; Vadez, 2014), very little is known about how plant-pathogen interactions affect root functioning and the hydraulic balance between roots and shoots. The difficulty of working with intact roots has meant that the analysis of the impact of viral infections on plant hydraulics has almost exclusively been limited to transpiration. Viral infections can modify stomatal development, density and conductance, leading to altered transpiration rates (Hall & Loomis, 1972; Lindsey, 1975; Culver & Padmanabhan, 2007; Erion & Riedell, 2012; Murray et al., 2016; Ertunç, 2020; Manacorda et al., 2021). Leaf osmolyte content and leaf relative water content (RWC) can also be altered by plant viruses (Xu et al., 2008; Mishra et al., 2022; Aguilar & Lozano-Duran, 2022; Mirzayeva et al., 2023). With respect to roots, El Aou-ouad et al. (2017) showed that infection with GLRaV-3 reduced whole-plant hydraulic conductivity by 14-40%. A recent study in grapevine showed a mild but significant decrease in Lp_r_ in response to GFLV infection (Jež-Krebelj et al., 2022). Therefore, viral infections clearly have an impact on plant water management, with the potential to alter not only the plant’s response to further abiotic stresses (Prasch & Sonnewald, 2013; Aguilar & Lozano-Duran, 2022) but also the capacity to impact on its competitive fitness. As recently pointed out, no universal physiological outcome exists for plant viral infections, and each pathosystem should be separately studied (Aguilar & Lozano-Duran, 2022).

Potyviridae constitute the largest plant virus family, accounting for 30 % of all known plant viruses. Its members have a broad host range that encompasses 57 plant families (Inoue-Nagata et al., 2022; Qin et al., 2023). Within the Potyviridae family, the Potyviruses (viruses in the genus *Potyvirus*), comprise 183 species (https://talk.ictvonline.org/taxonomy/) and represent the largest group of known plant-infecting RNA viruses (Revers & García, 2015; Wang, 2021) Potyviruses are flexuous, filamentous particles 650–950 nm in length and 11–20 nm in width with single-stranded, positive-sense RNA genomes 8–11 kilobases (kb) in size (Inoue-Nagata et al., 2022). Within the plant pathology community, considerable attention has been devoted to potyviruses, because the genus contains some of the more devastating plant viruses affecting wild plants and crop production worldwide (Gibbs et al., 2020), including potato virus Y (PVY), plum pox virus (PPV) and turnip mosaic virus (TuMV) (Qin et al., 2023). TuMV is one of the best studied potyviruses, partially due to its ability of infect dozens of species of economic importance (Walsh & Jenner, 2002) such as Brassicaceae plants, including the model plant *Arabidopsis thaliana*. Hence, the couple Arabidopsis-TuMV has become itself a plant pathosystem model (Nellist et al., 2022). Potyviruses trigger multifaceted pleiotropic effects in its hosts (Revers & García, 2015; Mäkinen, 2023) and TuMV has been shown to alter its host’s biochemical composition (Rodríguez et al., 2012; Prasch & Sonnewald, 2013; Manacorda et al., 2013), transcriptomic profile (Yang et al., 2007; Prasch & Sonnewald, 2013; Sánchez et al., 2015; Corrêa et al., 2020) and morphological and developmental parameters (Sánchez et al., 2015; Manacorda & Asurmendi, 2018).

To develop integrated pest management strategies, we need a deeper understanding of the host responses involved in the development of devastating potyvirus diseases (Mäkinen, 2023). An important aspect that warrants further attention is how viruses affect root functions (Vaisman et al., 2023), especially since there is evidence that viruses (including TuMV) migrate to the root soon after infection (Lunello et al., 2007) and can affect root morphology (Villordon & Clark, 2014; Chen et al., 2017; Vaisman et al., 2023) sometimes leading to devastating effects on the roots of cultivated crops (Bakaye et al., 2020; Peng et al., 2022).

We recently showed that TuMV infection significantly diminished stomatal conductance (g_s_), rosette transpiration rate and whole plant water consumption in Arabidopsis (Manacorda et al. 2021). We hypothesized that impaired transpiration would impact root hydraulics and the root-to-shoot water balance. In the present work, we provide a comprehensive analysis of TuMV-induced alterations in plant hydraulics at the whole-plant level, integrating measurements of root hydraulic conductivity, aquaporin contribution, and anatomical changes to elucidate the mechanisms underlying virus-mediated disruption of whole-plant water balance. These hydraulic alterations, along with their relationships to osmotic imbalances and the carbon economy, revealed a compromised plasticity under infection and reflected the pleiotropic physiological impact that TuMV exerted on the plant.

## Materials and methods

### Plant material and growth conditions

*Arabidopsis thaliana* ecotype Col-0 was used throughout this study. Hydroponic growth under a short-day photoperiod (10 h light/14 h dark) was described by Manacorda et al. (2021).

### Virus infection assays

TuMV-UK1 strain (GenBank accession AF169561;Sánchez et al., 1998) was propagated in infected *A. thaliana* plants. For each experiment, fresh infectious sap was prepared immediately before use. Viral infections were performed as described by Manacorda *et al*. (2021). To enable comparisons across experiments and parameters, all measurements were performed at the same stage of viral infection (12–13 days post-inoculation, DPI).

## Measurement of physiological parameters

### Relative water content

Leaf relative water content (RWC) was measured as previously described (Turner, 1981; Sade et al., 2015) using the 7th or 8th fully expanded rosette leaf. Turgid weight (TW), fresh weight (FW) and dry weight (DW) were recorded for each leaf and RWC values were calculated as 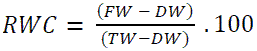.

### Determination of osmotic potential

The osmotic potential of the 7th or 8th fully expanded rosette leaf and roots was determined as described by Sutka et al. (2016). Sap osmolarity (mOsm kg⁻¹) was measured on a vapor pressure osmometer (Vapro 5520, Wescor, Logan, UT, USA) and used to calculate osmotic potential (MPa) via the van ‘t Hoff equation assuming 25 °C (R = 0.008314 MPa·L·mol⁻¹·K⁻¹).

### Whole root hydraulic conductivity

Measurements of root hydraulic conductance were performed as previously described (Vitali et al., 2021). The entire, intact root system of a freshly detopped plant was inserted into a 50 ml Falcon tube filled with BNS solution and was then placed in a Scholander-type pressure chamber (Model 4, BioControl, Argentina). The hypocotyl was carefully sealed to a glass capillary using a low-viscosity silicone dental impression material (A+Silicone, Densell) and the assembly was passed through the chamber lid. The roots were successively subjected to pressures (P) of 0.3, 0.4, and 0.2 MPa and the flow (J_v_) of the root exudate was measured. Hydraulic conductance (L_0_) was obtained from the slope of the plot of J_v_ versus P (*J*_*v*_ = *L*_0_. *P*). L_0_ values were then normalized by the dry weight of the entire root tissue to obtain Lp_r_ (Sutka et al., 2011), by the dry weight of the entire rosette tissue to obtain Lp_s_ (Maurel et al., 2010) or by the rosette projected area (PrA), to obtain Lp_A_ (Lo Gullo et al., 1998; Laur & Hacke, 2013). All root conductivity measurements were performed between ZT1 and ZT7 (Zeitgeber time; ZT0 = lights on).

### Aquaporin contribution to whole root hydraulic conductivity

To assess the contribution of AQPs to the hydraulic conductivity of the root, the effect of sodium azide, which blocks the acid-induced gating of AQPs, was studied as described by Sutka *et al*. (2011). Plants were equilibrated for 20 min at 0.3 MPa in a solution containing 5 mM KNO_3_, 2 mM MgSO_4_, 1 mM Ca(NO_3_)^2^ and 10 mM MES, pH = 6, supplemented with 20 mM KCl and exudate flow (J_v_) was recorded to determine Lp_r_. Then, 1 mM NaN₃ was added to the bathing solution and Jᵥ at 0.3 MPa was monitored for 30 min until a steady flow was reached. The maximum percentage inhibition of J_v_, and therefore of Lp_r_, was determined by fitting the kinetic curves with a negative exponential function.

## Determination of anatomical parameters

### Leaf and rosette area

To measure the total leaf area (TLA), all rosette leaves were excised, flattened without overlapping on a contrasting background, and scanned. To measure the rosette projected area (PrA), zenithal photographs were captured with a digital camera (Canon PowerShot SX50HS) against a contrasting background. TLA and PrA were quantified from the resulting images using ImageJ (https://imagej.net/ij/). In time-series experiments, photographs were taken at the same Zeitgeber time on successive days. Mean relative rosette growth rate (RGR) over a period Δt (between 0 and 12-13 DPI) was calculated as in De Vylder et al. (2012).

### Root anatomical measurements

To anatomically characterize the root and determine the histological characteristics of the vascular cylinder, freehand cross sections of the root were cut at different distances from the root apex and observed under either bright-field or epifluorescence microscopy with an Axioskop 2 microscope (Carl Zeiss, Jena, Germany) To determine the characteristics of the endodermis, freshly sectioned 10-mm-long root segments were incubated in 0.3% (w/v) Sudan IV in 70% ethanol for 1 h (Sutka et al., 2011) rinsed briefly in 70% ethanol, and then sectioned as described above.

### Aquaporin gene expression assays

Oligonucleotide primer sets for real-time (quantitative) PCR were designed using Primer BLAST software (Ye et al., 2012). They are listed in Supplementary Table S1. Details on the minimum information for publication of quantitative real-time PCR experiments (MIQE Guidelines: (Bustin et al., 2009, 2010)) including RNA extraction and cDNA protocols are listed in Supplementary Table S2. Primers were designed to target exon regions and for genes for which splice variants are predicted using ThaleMine, primers were designed to target all splice variants. For qPCR analysis, LinRegPCR program and the normalization method of Pfaffl et al., 2002 were used. Relative expression ratios and statistical analysis were calculated using fgStatistics software (Di Rienzo, 2009).

### Measurements of gas-exchange

Gas exchange measurements were performed with a LI-6400XT portable photosynthesis system (LiCor Inc., Lincoln, NE, USA). All measurements were carried out on intact plants at 12-13 DPI, using the youngest fully expanded leaf. When the leaf did not cover the entire chamber area (6 cm^2^), the measurements were corrected by the actual leaf area analysed, measured from digital photographs. The measuring chamber conditions were chosen to be similar to those experienced by the plants inside the growth chamber. The sample CO_2_ concentration was kept at 400 μmol mol^−1^, the leaf temperature at 22 ± 0.5°C and the relative humidity was set between 65 and 75%, with an air flow of 300 μmol s^−1^. Dark respiration was recorded after equilibration of the leaf in the dark, followed by at least 5 min of illumination with a photosynthetically active radiation of 150 μmol photons m^-2^ s^-1^ (90% red light/10% blue light, provided by the LI6400-02B LED light source) at the end of which a measurement of net photosynthesis and water vapor exchange was performed. Intrinsic water use efficiency was calculated as the ratio between net photosynthesis and stomatal conductance. All gas exchange measures were performed between ZT1 and ZT7.

### Statistical analysis

Aside from qPCR analyses, all statistical analyses were performed using R (RStudio Team, 2019; R Core Team, 2020). For all statistical analysis, significance was set as: NS = p > 0.05, * = p ≤ 0.05, ** = p ≤ 0.01, and *** = p ≤ 0.001. Normality assumptions were checked for all datasets prior to t-test or ANOVA analyses; those datasets that failed normality tests were analysed using nonparametric Wilcoxon or Kruskal – Wallis tests. For data presenting nested (hierarchical) or serial correlation (time-dependency) structure, linear mixed effects models (lme) and generalized least squares models (gls) were implemented using the nlme package of R (Pinheiro, Bates, DebRoy, & Sarkar, 2020), used as suggested by (Pinheiro & Bates, 2000) for constructing, validating, comparing and selecting models. Multiple comparison tests were performed using the emmeans package (Lenth, 2019). Effect size (Cohen’s d) was calculated based on (Lakens, 2013).

## Results

### Impact of TuMV on plant growth and root anatomical parameters

We evaluated the impact of TuMV infection on biomass and growth parameters using a hydroponic *Arabidopsis thaliana* (Col-0)–TuMV-UK1 pathosystem grown under short-day conditions (Fig. 1, Supplementary Table S3). Compared to mock-inoculated plants, TuMV infected plants exhibited significantly reduced dry weight in both rosettes and roots (by 35% and 44%, respectively; p < 0.001) and biomass partition (dry weight shoot/dry weight root = DWsr) was similar between treatments (p> 0.05; % change = –3.3%). Additionally, TuMV infection reduced rosette projected area (PrA) by 58% (Fig. 1B), and PrA was highly correlated with total leaf area (Supplementary Fig. S1). The impact on PrA and shoot biomass was not proportional, as specific leaf area (SLA) decreased by 30% under infection (Fig. 1C).

**Figure 1.**
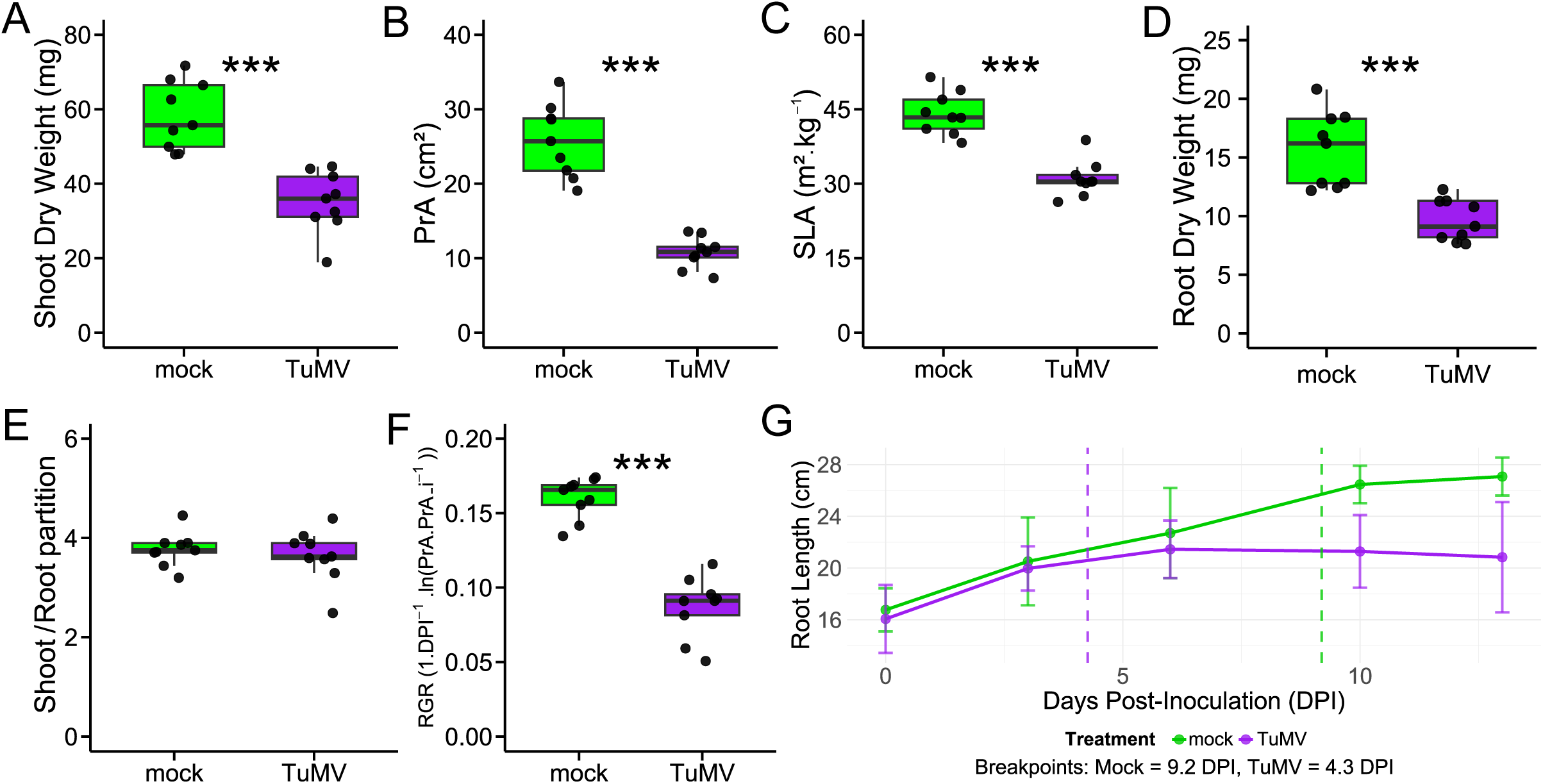
TuMV alters shoot and root biometric and growth parameters. (A) Shoot (rosette) dry weight; (B) projected rosette area (PrA); (C) specific leaf area (SLA); (D) root dry weight; (E) dry biomass partition (shoot-to-root ratio); (F) rosette relative growth rate (RGR), calculated as 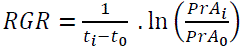; (G) breakpoint analysis showing apical root growth dynamics (dashed lines = breakpoints). Parameters were measured in healthy, mock-inoculated plants (green) and in TuMV-infected plants (purple). (A-F) At least three independent experiments were performed and results from one representative experiment are shown. n = 9 for each treatment. Each dot represents an individual plant, and the boxes display the median (center line) and interquartile range. *** = p ≤ 0.001. (G) Two similar independent experiments were combined weighing treatment means for each DPI. 8 ≤ n ≤ 15 per treatment and experiment. Dashed lines represent breakpoint time and error bars the confidence interval.

Analysis of rosette and root growth dynamics further revealed the negative impact of TuMV infection. The virus caused a decrease in rosette relative growth rate (RGR) of 45% between 0 and 13 DPI (p value < 0.001; Fig. 1F). In the same line, we observed a considerable arrest in primary root length, where breakpoint analysis on root length growth dynamics showed that for healthy plants, root length reached a breakpoint at 9.2 ± 1,3 DPI, whereas infected plants anticipated the breakpoint by 5 days, at 4.3 ± 0.1 DPI (Fig. 1G). Segmented regression plateaus indicate that apical root elongation declined to near-zero beyond 3–5 DPI in infected plants (Fig. 1G; Supplementary Table S4). We note that this inference relies on length plateaus rather than direct assays of meristem activity or root tip viability, which were not performed here. Altogether, these results highlight how viral infection affects both aboveground and belowground organs, impairing overall plant growth in agreement with results observed in aerial parts of soil-grown plants (Manacorda & Asurmendi, 2018).

To assess the impact of the viral infection on root anatomy, we first analysed cross sections of the absorption (root hair) zone of the root, taken 10 mm from the root tip, and focused on the radial organization of tissues and characteristics of the endodermis. Sudan IV staining was used to assess suberin deposition in the endodermis, and cross sections were also observed under a confocal microscope to primarily visualize autofluorescence of suberin and lignin in the endodermis and xylem respectively. No differences were observed in the level of endodermis suberization between mock– and TuMV-inoculated plants (Fig. 2 A-F). A small, non-significant diameter reduction was detected for roots, the vascular cylinder, and the xylem vessels (Fig. 2G). Accordingly, in mock-inoculated plants, the cortex in the absorption zone of roots comprised, in almost all sections sampled, two layers of parenchymatous cells, whereas in the same region of TuMV-infected roots there was a considerable proportion of cortex with only one cell layer (odds ratio = 7; p = 0,01; Fig. 2H).

**Figure 2.**
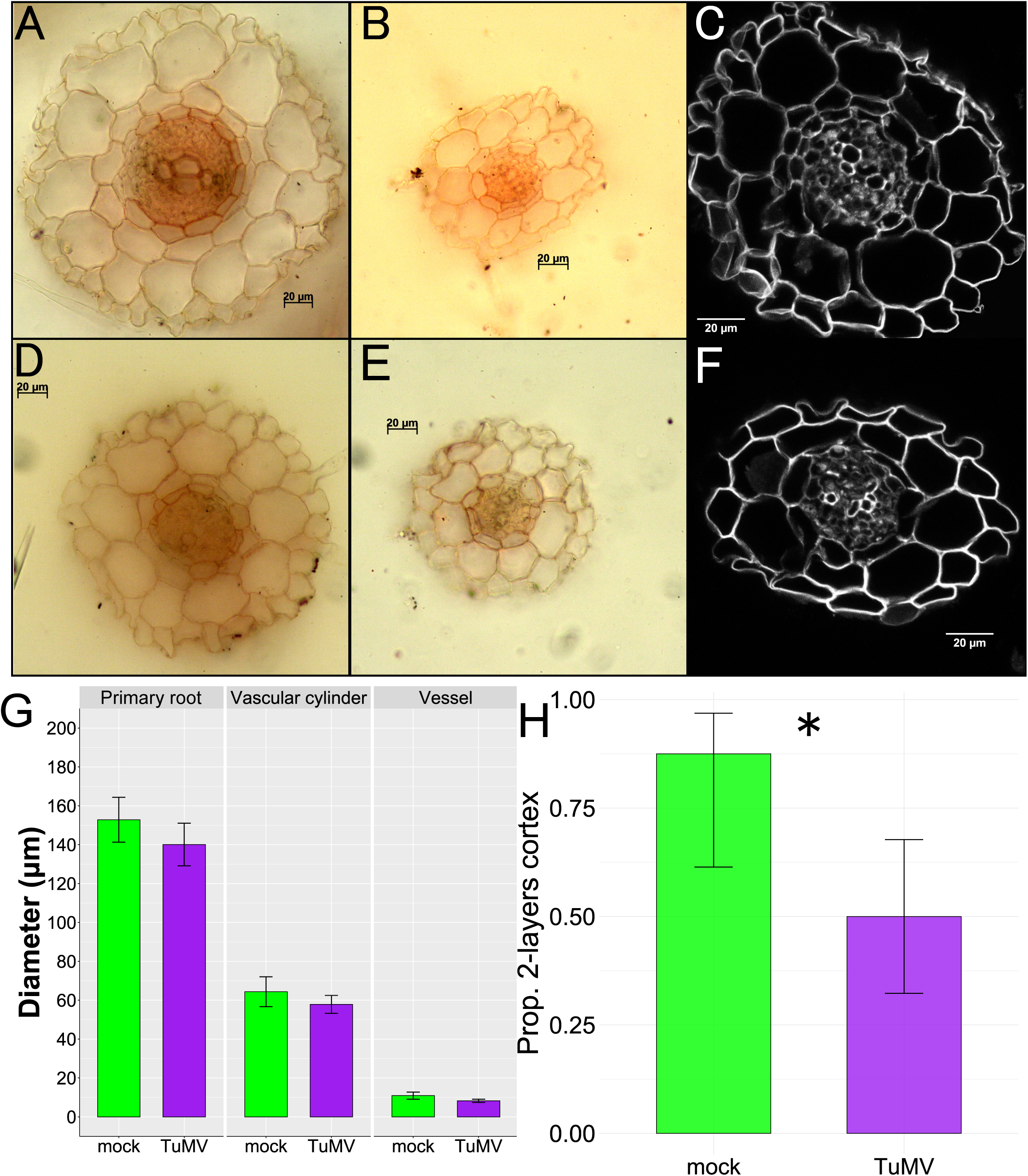
Root histology reveals impaired cortical cell growth under infection. Transverse root sections of (A-C) mock and (D-F) TuMV-inoculated plants stained with Sudan IV to identify suberin (A, B, C, E) or visualized by autofluorescence to identify lignin (C, F). (G) Quantification of primary root, vascular cylinder or vessel diameters. (H) Proportion of root sections with two cortical layers in mock and TuMV-infected plants. Shown are means ± 95% CI. n = 4 plants/treatment, 3-10 sections/plant. p < 0.05. Two independent experiments were performed, results from one representative experiment are shown.

### TuMV impairs carbon assimilation and alters gas-exchange balance

Given the marked reduction in biomass accumulation and rosette expansion described above, we next asked whether photosynthetic carbon gain and leaf gas exchange were also affected. To address this, we measured leaf gas exchange under the prevailing growth light conditions. TuMV infection significantly altered most gas-exchange parameters, whereas dark respiration (*R*_d_) remained unchanged between treatments (Fig. 3A). Infected plants exhibited significantly lower CO₂ assimilation rate (A; Fig. 3B) and g_s_ (Fig. 3C), with the virus having a more marked effect on conductance than on CO_2_ assimilation, which led to a higher intercellular CO_2_ concentration (Ci) and intrinsic water use efficiency (WUE) in infected plants compared to mock-inoculated ones (Fig. 3E and 3F). Integrating these measurements over a full diurnal cycle, the calculated carbon balance, estimated from the gas-exchange data, was 59% lower in infected plants than in controls.

**Figure 3.**
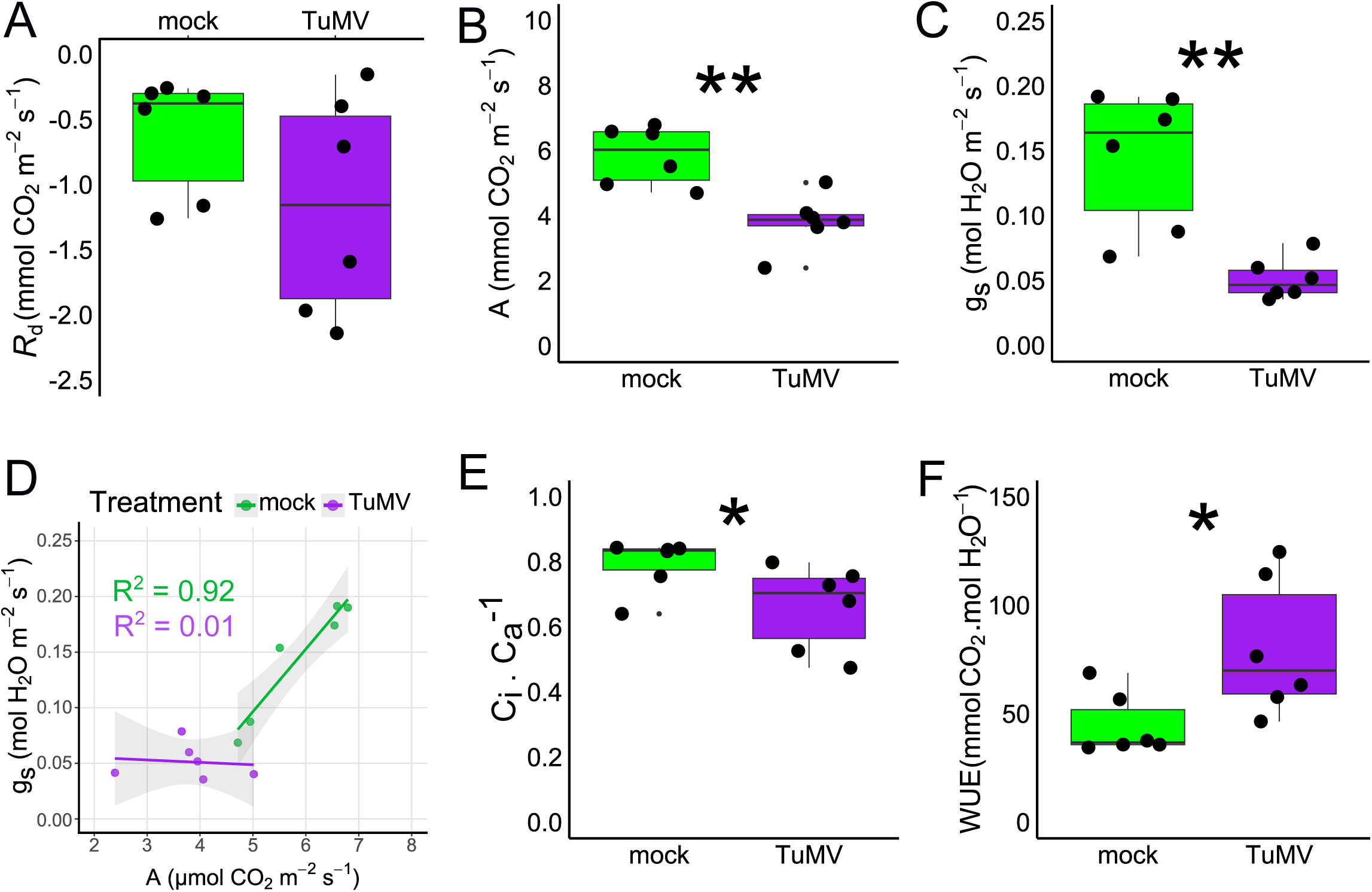
TuMV infection led to a disrupted carbon and water economy. (A) Apparent rate of respiration in the dark (Rd); (B) net CO_2_ assimilation (A); (C) stomatal conductance (g_s_); (D) correlation between g_s_ and A (data fitted to a linear equation); (E) ratio of intercellular to ambient CO_2_ (Ci/Ca); (F) intrinsic water use efficiency (WUE = A/g_s_). Each dot represents an individual plant, and the boxes display the median (center line) and interquartile range. n = 6. Two independent experiments were performed. Results from one representative experiment are shown.

### Viral infection reduces root hydraulic conductance but preserves rosette water-supply sufficiency

Building on our observation that TuMV impairs root development and reduces plant water loss and intrinsic water-use efficiency (Manacorda et al., 2021; Fig. 3), we next asked how the root system adjusts its hydraulic properties under infection. We therefore quantified root hydraulic conductance in TuMV-infected and mock-inoculated plants using the pressure-chamber method (Sutka et al., 2011; Fig. 4). Root hydraulic conductance (L_0_) was significantly lower in infected plants (Fig. 4A). Because infected plants also displayed reduced biomass and leaf area, we further expressed L_0_ with three complementary normalizations. When normalized by root dry weight (Lp_r_), root hydraulic conductivity reflects the intrinsic water transport efficiency per unit root biomass, i.e. the functional capacity of the root system itself. Lp_r_ was significantly reduced in TuMV-infected plants relative to controls (Fig. 4B), indicating a compromised root water transport capacity. Normalizing L_0_ by shoot dry weight (Lp_s_) revealed a smaller but still significant decrease under infection (Fig. 4C). In contrast, L_0_ normalized by leaf area (Lp_A_) did not differ between treatments under our growth-chamber conditions (PAR 150 µmol m−2 s−1, 22°C, 65–75% RH; Fig. 4D). Because Lp_A_ relates to the sufficiency of the roots to supply water to the transpiring leaf area (Laur & Hacke, 2013), this stability suggests that, despite the strong reduction in root biomass and Lp_r_, infected plants maintain a root–shoot hydraulic balance that is adequate for the lower transpirational demand of their reduced rosette.

**Figure 4.**
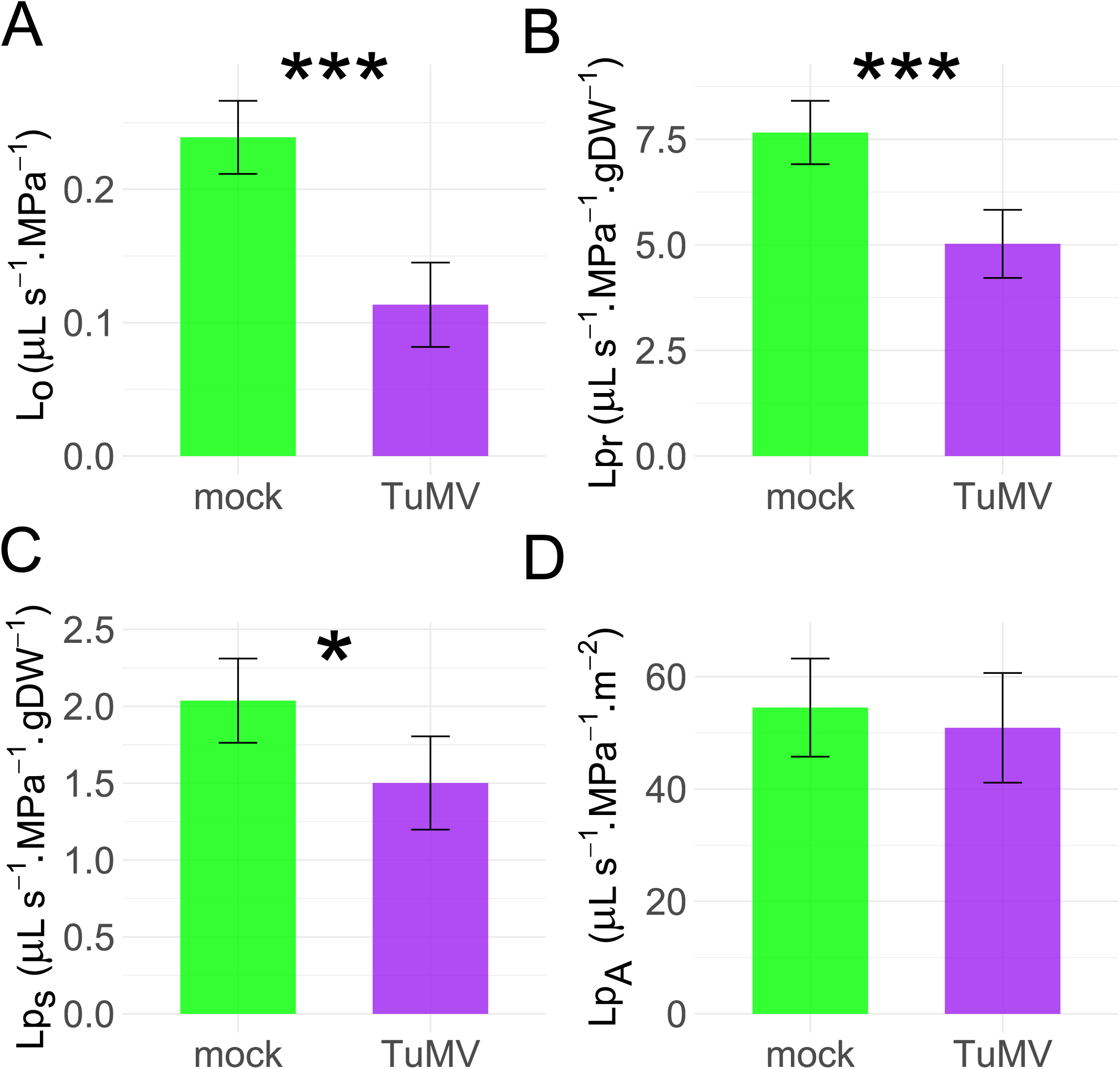
TuMV infection alters root hydraulic parameters. (A) L_0_, root hydraulic conductance, (B) Lp_r_, root hydraulic conductance normalized by root dry weight; (C) Lp_s_, root hydraulic conductance normalized by rosette dry weight (D) Lp_A_, root hydraulic conductance normalized by rosette projected area. Data are from one representative experiment (n = 12-16 plants per treatment) of three independent experiments that showed similar results. Means with error bars representing 95% confidence intervals are shown. * = p ≤ 0.05, *** = p ≤ 0.001.

### TuMV downregulates the expression of aquaporins without affecting their relative flow contribution

As it is well known that aquaporins are largely influenced by the cell-to-cell pathway, we analysed whether the virus affected their relative contribution to root total water flow. We quantified the degree of inhibition of the flow, J_v_, of the xylem exudate of excised roots in a pressure chamber at 0.3 MPa in the presence of sodium azide (NaN_3_). By blocking the cytochrome-mediated mitochondrial respiration, NaN_3_ induces an intracellular acidosis which leads to H^+^-dependent gating of plasma membrane AQPs (Tournaire-Roux et al., 2003; Boursiac et al., 2022). In the presence of NaN_3_, we observed a strong inhibition of J_v_ in both treatments, indicating that the AQPs contribute to approximately 70% of the root water transport (Fig. 5A), a result that is consistent with the AQPs as an important component in the modulation of J_v_ (Tournaire-Roux et al., 2003; Sutka et al., 2011) The fact that, in comparison with mock-infected plants, TuMV-infected plants maintained an unchanged percentage inhibition by NaN_3_ of root J_v_ indicates that the virus did not cause a significant change in the proportional contribution of the cell-to-cell and the apoplastic pathway to the overall water transport.

**Figure 5.**
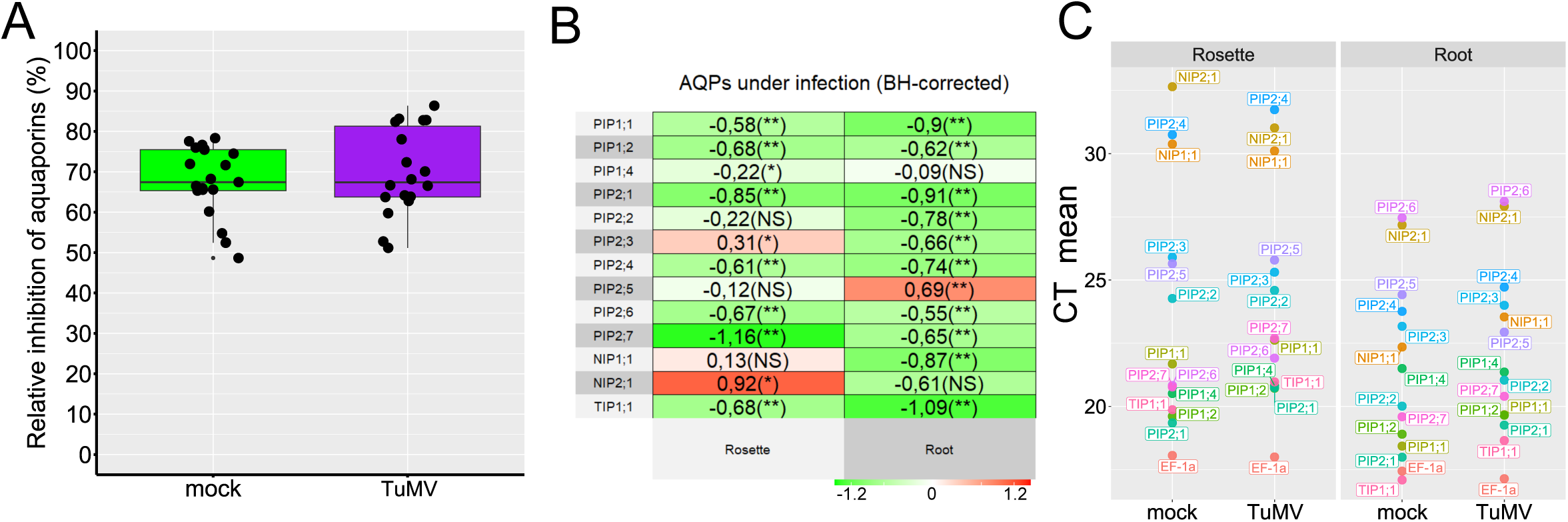
Infection with TuMV led to downregulation of mRNAs encoding water transport–related aquaporins. (A) Percentage inhibition of Lp_r_ by sodium azide, showing the contribution of aquaporins to cellular water transport pathways. Two independent experiments which yielded similar results were combined using a linear mixed-effects model with experiment as a random factor. n = 21 for mock-inoculated plants and n = 18 for TuMV-infected plants. (B) Heatmap of relative quantification of AQPs transcript abundance by qRT-PCR (log2 fold change). P-values are corrected for multiple comparisons by the Benjamini-Hochberg algorithm. n = 6 per group. Green, negative values = downregulation by TuMV; Red, positive values = upregulation by TuMV. (C) Expression level of 13 AQPs relevant for water transport and stress response. CT mean values indicate relative transcript abundance for each organ and treatment. Lower CT values are indicative of higher expression levels. The CT value of the gene EF-1a was included as a reference.

We next evaluated the gene expression of main water-transporting AQPs in roots and shoots of plants exposed to the virus. including NIP2;1 and PIP2;5, which are known to be induced under abiotic stress conditions. Under TuMV infection, there was a widespread and significant downregulation of key water-transporting AQPs in root tissue. In rosette tissue the downward pattern was present but less homogeneous in nature (Fig. 5B). Notably, PIP2;5 and NIP2;1 exhibited a distinct organ-specific induction, being upregulated by TuMV in roots and rosettes, respectively, from comparatively low basal levels in those organs. We then compared the relative baseline abundance of major AQP transcripts between organs using C_T_ values from qPCR analyses as a proxy for mRNA levels (Fig. 5C). This approach allowed the identification of tissue-specific expression differences: whereas several AQPs were expressed at similar levels in both roots and rosettes, others such as NIPs and PIP2;4 showed lower expression levels in the rosette. PIP2;6, in contrast, was markedly more abundant in rosettes.

### TuMV promotes osmotic adjustment and increased tissue hydration in leaves without altering root osmotic potential

To assess plant water status under TuMV infection, we quantified leaf water content, relative water content (RWC) and osmotic potential in leaves and roots. In line with the observed reductions in g_s_ and the maintained sufficiency at the whole-plant level, infected plants displayed a significant decrease in leaf water content compared with mock-inoculated plants (Fig. 6A). In contrast, leaf RWC was higher in infected plants (Fig. 6B), indicating that, relative to their fully hydrated state, infected leaves maintained or even improved their hydration status. This was associated with a more negative leaf osmotic potential (Fig. 6C). consistent with osmotic adjustment, whereas root osmotic potential did not differ between treatments (Fig. 6C).

**Figure 6.**
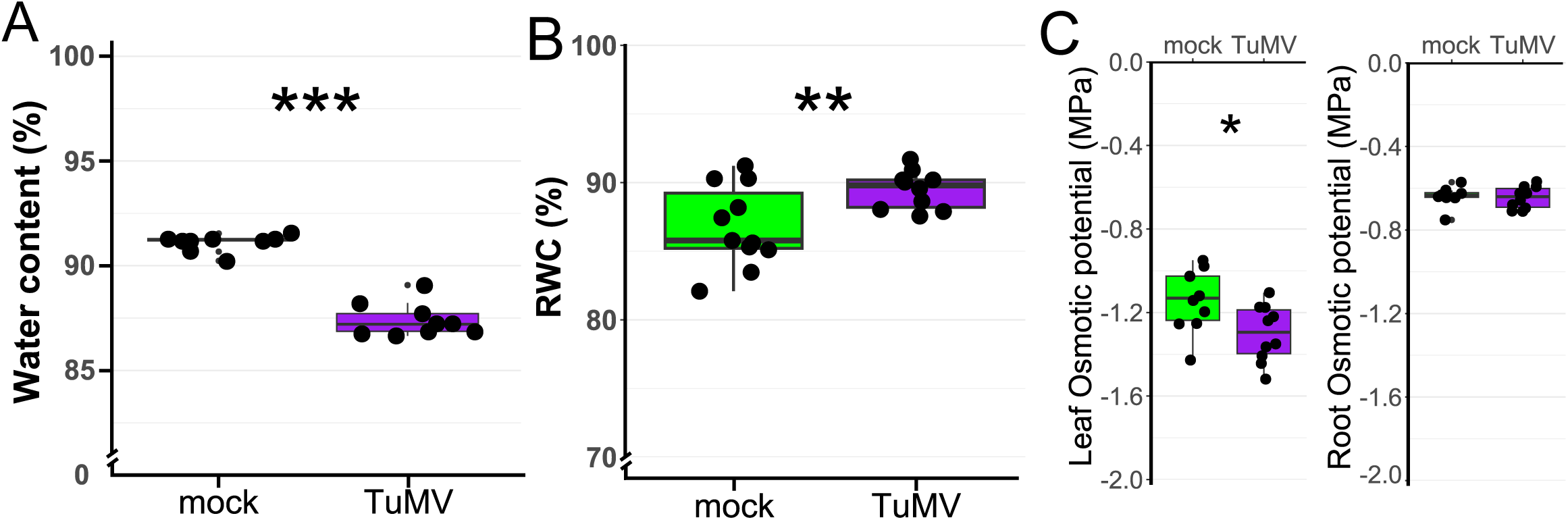
TuMV infection induces osmotic adjustment and enhanced water retention. (A) Percent of water content; (B) relative water content (RWC) of rosette leaves; (C) leaf and root osmotic potential. Three independent experiments were performed which showed similar results. One representative experiment is shown. n = 9-11. Each dot represents an individual plant, and the boxes display the median (center line) and interquartile range. * = p ≤ 0.05, ** = p ≤ 0.01, *** = p ≤ 0.001.

### Multivariate analysis identifies coordinated physiological shifts under TuMV infection

To evaluate the physiological response to TuMV infection from an integrated perspective, we applied multivariate inferential and descriptive statistics to a comprehensive set of biometric, anatomical, and hydraulic traits. PERMANOVA based on Gower distance revealed a highly significant multivariate difference between treatments (p < 0.001). Principal component analysis (PCA) on standardized variables confirmed a clear separation along the first plane, which explained the majority of total variance (85.9%; Fig. 7). Mock-inoculated plants clustered in the region associated with high values of specific leaf area (SLA), root hydraulic conductance (L₀), and biomass-related traits (projected rosette area, shoot and root dry weight), whereas TuMV-infected plants consistently exhibited lower values for these same variables. Notably, traits related to plant physiology and water transport, particularly L₀, SLA, Lp_r_, and Lp_s_, showed the highest loadings on the treatment-separating direction, while structural or partitioning traits such as biomass allocation (DWsr) and area-based hydraulic conductance (Lp_A_) contributed minimally to the observed divergence, indicating they were largely unaffected by TuMV infection. This pattern was further corroborated by hierarchical clustering on principal components (HCPC), which grouped individuals into two distinct clusters that aligned almost perfectly with infection status: the link between the cluster variable (treatment) and the categorical variables was highly significant (p = 1.5e-06, chi-square test). 100% of mock plants and 90% of TuMV-infected plants were correctly assigned to their respective clusters. The automatic report showed that, in descendent order, the variables that correlated the most with healthy plants were Lp_r_, Lo, Lp_s_, SLA and PrA (Vaissie et al., 2015). Together, these results demonstrate that TuMV infection induces a coordinated, system-wide downregulation of physiological performance, rather than an isolated or purely anatomical effect.

**Figure 7.**
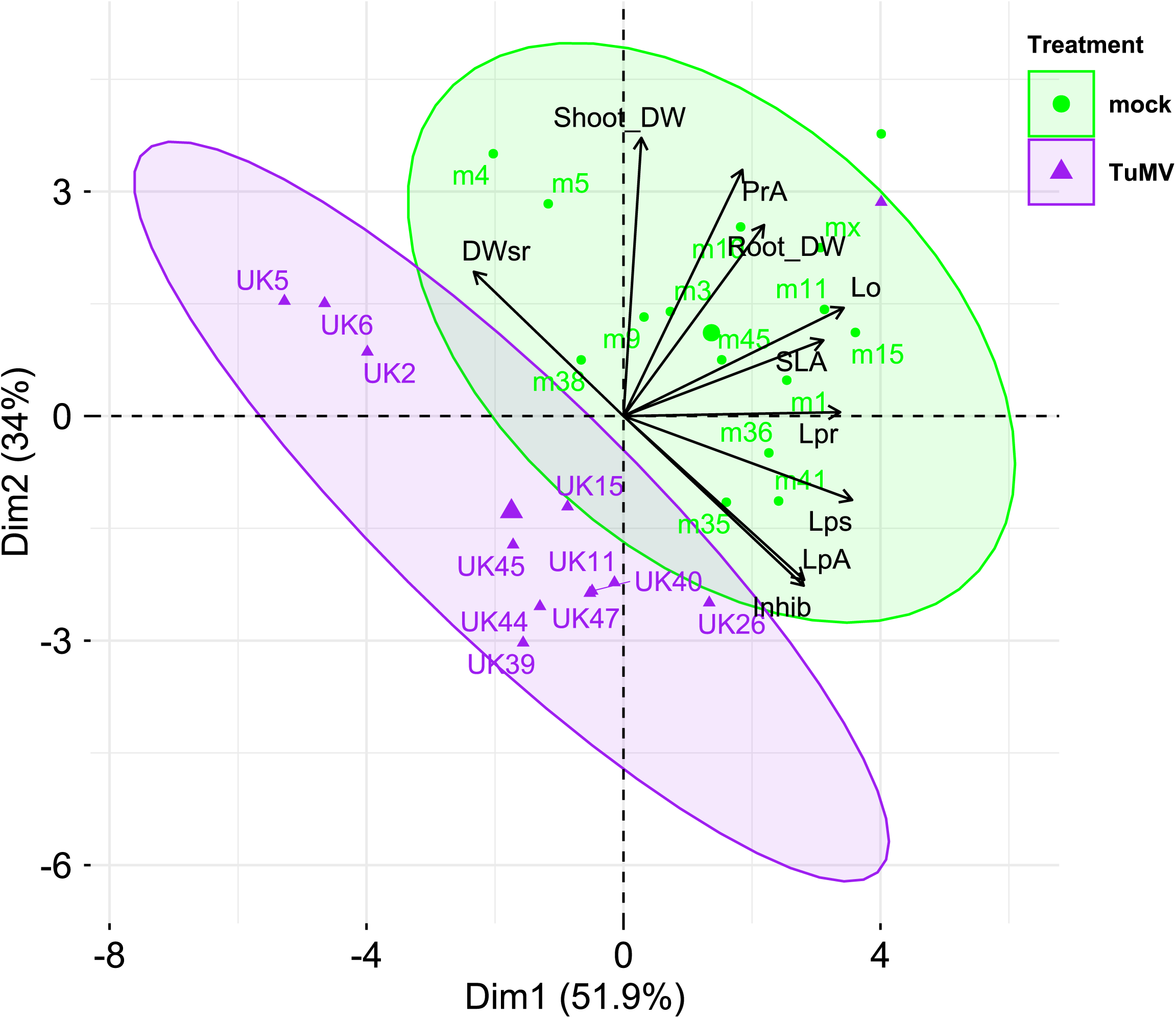
Principal Component Analysis (PCA) biplot of standardized biometric and physiological traits. The first two principal components explain 86% of the total variance. Symbols represent individual plants, coloured by treatment (green: mock; purple: TuMV). Ellipses denote 95% confidence ellipses around group centroids.

## Discussion

In this study, we provide an integrated, whole-plant physiological analysis of how infection by *Turnip mosaic virus* (TuMV) affects water management in *Arabidopsis thaliana*. Although hydroponic systems are widely used in plant nutrition and abiotic stress research, to our knowledge, they have only rarely been combined with viral pathosystems, and at the same time, hydraulic responses to virus infection have been examined in detail only in a few woody hosts (El Aou-ouad et al., 2017; Jež-Krebelj et al., 2022). By exploiting a hydroponic Arabidopsis–TuMV system that provides full access to intact roots, we integrate biometric, anatomical, hydraulic, osmotic, and photosynthetic measurements, and we demonstrate that TuMV infection triggers a marked reconfiguration that profoundly alters plant hydraulics, consistent with the severe growth arrest observed. This whole-plant approach moves beyond the well-documented effects of viruses on stomata and photosynthesis to reveal critical, previously underappreciated impacts on root function and root–shoot hydraulic integration.

Our work confirms and extends previous observations that TuMV infection causes severe growth and developmental impairment (Manacorda & Asurmendi, 2018; Fig. 1). The significant reduction in specific leaf area (SLA) indicates a marked shift in leaf structural traits. Allometric analyses of shoot-to-root biomass partitioning (Fig. 1E) showed no significant difference between treatments, indicating that the proportional reduction in root and shoot biomass is coordinated, further underscoring the integrated nature of the growth arrest. Our growth analysis reveals a dramatic and premature cessation of primary root elongation, with a breakpoint occurring five days earlier in infected plants (Fig. 1G). This arrest, which coincides with the reported timing of TuMV arrival in roots (Lunello et al., 2007), would be expected to compromise the plant’s ability to explore the soil for water and nutrients, a key trait for drought avoidance (Schenk & Jackson, 2002). This is consistent with our previous report that showed that TuMV had diminished tolerance to drought stress when competing with healthy plants (Manacorda et al., 2021). Our qualitative scoring revealed an increased frequency of roots with a single cortical layer (Fig. 2), further expanding the developmental disruption phenotype observed in the rosette (Manacorda & Asurmendi, 2018).

A small reduction in the diameters of primary roots, vascular cylinders, and xylem vessels was observed in TuMV-infected plants at 12 days post-inoculation (DPI). Although these changes did not reach statistical significance, they are consistent with the significant reduction in the average number of cortical cell layers (Fig. 2H) and fit within the broader pattern of TuMV-induced root growth impairment. This aligns with the more pronounced histological alterations reported by López-González et al., (2021) in mature inflorescence stems of TuMV-infected Arabidopsis at 20 DPI, suggesting that the extent of viral effects on plant anatomy may vary with tissue type, developmental stage, and time post-infection. Such systemic developmental disruption could stem from viral interference with hormone transport or signalling, which are known targets of viral manipulation (Vitti et al., 2013; Vaisman et al., 2023).

Consistent with the systemic reduction in growth rate, gas-exchange measurements showed that TuMV-infected plants operated with persistently lower g_s_ and CO₂ assimilation rates than controls. This sustained cost in daily carbon gain provides a mechanistic explanation for the strong reduction in biomass accumulation in infected plants. Additionally, because the decline in gₛ caused by infection was proportionally larger than the decline in assimilation, intrinsic water-use efficiency increased (Fig. 3F). In mock-inoculated plants, as expected, A was tightly coupled to g_s_ (R² = 0.92; Fig. 3D), whereas this relationship was essentially lost in TuMV-infected plants (R² = 0.01; Fig. 3D). This divergence is consistent with additional non-stomatal limitations on photosynthesis, which could reflect direct damage to the photosynthetic machinery or virus-induced metabolic reprogramming, as previously reported for TuMV in Brassicaceae (Guo et al., 2005). Thus, the measured increase in intrinsic WUE likely reflects a pathological decoupling of carbon and water fluxes rather than an adaptive strategy aimed solely at conserving water, in contrast to the patterns broadly observed in drought-stressed leaves (Grimmer et al., 2012; Hummel et al., 2010). The loss of functional coordination between gas-exchange parameters resulted in a 59% collapse of diurnal carbon balance, providing a direct physiological explanation for the observed growth retardation. This breakdown in leaf-level integration signals a systemic failure to maintain physiological homeostasis under viral stress an alteration that prompted us to examine root physiology in detail.

The impact of viral infection on root development is coupled with a significant alteration in root physiology. We found that TuMV infection reduces root hydraulic conductivity (Lp_r_; Fig. 4). This marked reduction in Lp_r_ reveals a lower intrinsic capacity of the root system to transport water per unit biomass, a clear hydraulic limitation imposed by the virus at the root level. Importantly, this reduction in Lp_r_ occurs in parallel with a strong decrease in g_s_, which is consistent with the view that plants regulate water fluxes mainly through coordinated downregulation of root and shoot hydraulic conductance and stomatal conductance along the soil–plant–atmosphere continuum (Sade et al., 2009; Maurel et al., 2010; Vandeleur et al., 2014; Moshelion et al., 2015; Maurel et al., 2016). Even under hydroponic conditions with non-limiting water supply, roots thus emerge as active regulators of whole-plant hydraulics rather than as passive conduits. When hydraulic conductance is normalized by shoot dry weight (Lp_s_) or, more importantly, by projected leaf area (Lp_A_), the difference between treatments diminishes or disappears, indicative of maintained hydraulic sufficiency per unit leaf area under the tested conditions. However, the apparent stability of Lp_A_ largely reflects the reduced g_s_ and leaf area of infected plants. If g_s_ were restored to healthy levels without a concomitant increase in Lp_r_, the diminished root system would likely be unable to sustain water demand, leading to rapid hydraulic imbalance. Therefore, stable Lp_A_ values under infection reinforces the concept of a tight coupling between root hydraulic capacity and shoot transpiration and shows that such coordination is functionally conserved under severe biotic stress.

This resulting hydraulic balance was further explored through our analysis of AQP function and gene expression. We first assessed the functional contribution of AQPs to root water transport using NaN₃ inhibition. Despite TuMV infection, the relative AQP contribution to total root hydraulic conductance remained unchanged at ∼70% (Fig. 5A), indicating that AQPs continue to represent the principal route for water flow in the presence of the virus. In roots, and in line with the ∼25% reduction in Lp_r_, TuMV infection led to a widespread downregulation of AQP transcripts (Fig. 5B). This pattern suggests that the plant does not reconfigure the functional partitioning among AQP-mediated, plasmodesmatal (symplastic) and apoplastic pathways; rather, it downscales overall capacity while preserving their relative contributions. The maintenance of this AQP contribution under infection underscores a flexible, well-regulated hydraulic control point. Within the symplastic pathway of water flow, plasmodesmata constitute another checkpoint to consider. Given that viruses such as TuMV interact with plasmodesmata for their movement (Wang, 2021) and that virus–host interactions can involve complex responses affecting plasmodesmal function (Huang et al., 2023), alterations in intercellular transport pathways may contribute to the observed Lp_r_ reduction without altering pathway partitioning.

At the transcriptional level, most AQPs followed this global trend towards downregulation. The only consistent exceptions were the stress-responsive NIP2;1 in rosettes and PIP2;5 in roots, which were upregulated upon infection. However, their transcript levels remained low compared with other AQPs (Fig. 5C), suggesting that they are unlikely to counterbalance the overall downregulation of water transport capacity. Both isoforms have been associated with metabolic or abiotic stress responses (Jang et al., 2004; Beamer et al., 2021; Jiang et al., 2023, 2025), pointing to a more specific role in stress adjustment rather than in bulk hydraulic regulation. Further investigations are needed to assess AQP activity in order to obtain a clearer picture of their role under viral infection.

The maintenance of hydraulic sufficiency (Lp_A_) is accompanied by a shoot-localized osmotic adjustment: infected plants show a more negative leaf osmotic potential together with higher RWC (Fig. 6), consistent with solute accumulation that helps preserve leaf hydration and tissue function when water influx is reduced. Notably, this adjustment appears shoot-specific while root osmotic potential remains unchanged. Together with the infection-induced downscaling of gas exchange described above, this shoot-localized osmotic adjustment provides a coherent physiological explanation for the observed growth penalties under infection.

Our results show that the loss of physiological plasticity caused by TuMV infection extends beyond the shoot. From a hydraulic perspective, it is also evident in the constitutive downregulation of root aquaporins and the associated reduction in Lp_r_. Under normal conditions, these parameters are dynamically regulated to fine-tune water uptake in response to environmental fluctuations. However, TuMV infection appears to lock the root hydraulic system into a low-conductivity state, uncoupling it from the potential for dynamic adjustment, which is critical for stress responses (Maurel et al., 2010). Consequently, the infected plant is not merely smaller or less efficient; it is fundamentally less adaptable, rendering it highly vulnerable to any additional environmental challenge (Prasch & Sonnewald, 2013; Manacorda et al., 2021).

The multivariate integration of anatomical, biometric, and hydraulic traits (Fig. 7) reveals a striking pattern: physiological variables are strong drivers of phenotypic divergence between mock and TuMV-infected plants, especially those related to water transport (L₀, Lp_r_) and leaf economics (SLA), whereas traits reflecting structural allocation (e.g., shoot-to-root biomass ratio) or hydraulic sufficiency per unit leaf area (Lp_A_) remain largely conserved. This finding underscores that the core impact of TuMV goes beyond a reduction in size, provoking a profound reprogramming of the plant’s functional physiology. In other words, the virus does not simply make a “smaller plant”; but a physiologically distinct organism with compromised plasticity, reduced carbon gain, and locked hydraulic capacity. This systemic functional shift, detectable even in hydroponic, non-drought conditions, highlights the value of whole-plant physiological phenotyping as a critical lens for understanding plant–virus interactions beyond symptomatology or growth metrics alone.

In conclusion, Arabidopsis Col-0 plants with TuMV-UK1 infection orchestrates a systemic physiological response that prioritizes short-term survival (maintaining leaf hydration via osmotic adjustment and reducing water loss via stomatal closure) at the expense of long-term growth and fitness. The plant achieves a new, suboptimal equilibrium where a stunted root system with reduced absolute hydraulic conductivity is nonetheless sufficient (Lp_A_) to supply a stunted shoot with its reduced water demands. From an ecological perspective, this hydraulic “downsizing” makes infected plants poor competitors for water in mixed stands, as their healthy neighbours with higher Lp_r_ will deplete soil moisture more rapidly (Casper & Jackson, 1997), which can help explain previous results (Manacorda et al., 2021). Our holistic approach, linking viral infection to root hydraulics, shoot physiology, and whole-plant carbon economy, provides a crucial framework for understanding the complex and often devastating physiological consequences of plant virus infections. An integrative, multi-level view of plant–virus interactions helps avoid one-sided interpretations and advances our understanding of infection dynamics (Pollari et al., 2022). In this framework, physiological analyses provide the functional context that anchors molecular changes to whole-plant and ecological performance.

## Supplementary data

**Supplementary Table 1:** List of primers used for qRT-PCR.

**Supplementary Table 2:** Experimental conditions used for messenger RNA quantification by qRT-PCR assays based on MIQE requirements (Bustin *et al*., 2009; Bustin *et al*., 2010).

**Supplementary Table 3:** Summary statistics for biomass and growth variables following TuMV infection.

**Supplementary Table 4:** Premature cessation of apical root growth in infected plants.

**Supplementary Figure 1:** Correlation between total leaf area and projected resette area

## Acknowledgements

We thank Dr. Flora Sánchez and Dr. Fernando Ponz for the kind gift of TuMV virus UK1 strain. We also thank Marina Recchi for expert technical assistance. AI was used for help with English language editing.

## Author contributions

All authors conceived and designed research. CAM, PDC and MS performed the experiments. CAM, PDC and IB prepared the manuscript with a final editing by SA, IB and GA. SA, GA and IB. obtained funding; all authors discussed the results and read and approved the manuscript. None of the authors have a conflict of interest to disclose.

## Funding

This work was supported by grants from the Agencia Nacional de Promoción Científica y Tecnológica, Argentina [PICT 2020-01438] to G.A. and [PICT 2021-697] to A. S.; from Instituto Nacional de Tecnología Agropecuaria [2023-PD-L01-I084] to A. S.; from Consejo Nacional de Investigaciones Científicas y Técnicas, Argentina [PIP 11220200101935CO] to I.B. and from University of Buenos Aires, Argentina [UBACyT 20020220100162BA] to I.B.

## Figure Legends

**Figure Supplementary 1.** Correlation between total leaf area and projected rosette area. Total leaf area (TLA) and rosette projected area (PrA) are highly correlated for both treatments. The percentage of superimposed leaves range from 16 to 26 %, without significant differences between treatments. Area is given in pixels (px). Data taken from plant 13 DPI (n = 6-8 plants per treatment).

## Notes

### Competing Interest Statement

The authors have declared no competing interest.

## References

Aguilar E., Cutrona C., Del Toro F.J., Vallarino J.G., Osorio S., Pérez-Bueno M.L., … Tenllado F. (2017) Virulence determines beneficial trade-offs in the response of virus-infected plants to drought via induction of salicylic acid. Plant, Cell & Environment 40, 2909–2930, 10.1111/pce.13028.

Aguilar E. & Lozano-Duran R. (2022) Plant viruses as probes to engineer tolerance to abiotic stress in crops. Stress Biology 2, 20, 10.1007/s44154-022-00043-4.

Aroca R., Porcel R., & Ruiz-Lozano J.M. (2012) Regulation of root water uptake under abiotic stress conditions. Journal of Experimental Botany 63, 43–57, 10.1093/jxb/err266.

Bakaye D., Amadou H.B., Sognan D., Salimatou S., Adounigna K., Rokiatou F., … Linda K. (2020) Inhibition of root growth as mode of action of two rice yellow mottle virus pathotypes isolated in Mali. African Journal of Agricultural Research 16, 1148–1154, 10.5897/AJAR2020.14929.

Boursiac Y., Protto V., Rishmawi L., & Maurel C. (2022) Experimental and conceptual approaches to root water transport. Plant and Soil 478, 349–370, 10.1007/s11104-022-05427-z.

Boyer J.S. (1985) Water Transport. Annual Review of Plant Physiology 36, 473–516, 10.1146/annurev.pp.36.060185.002353.

Buckley T.N. (2019) How do stomata respond to water status? New Phytologist 224, 21–36, 10.1111/nph.15899.

Bustin S. a, Beaulieu J.-F., Huggett J., Jaggi R., Kibenge F.S.B., Olsvik P. a, … Toegel S. (2010) MIQE précis: Practical implementation of minimum standard guidelines for fluorescence-based quantitative real-time PCR experiments. BMC molecular biology 11, 74, 10.1186/1471-2199-11-74.

Bustin S. a, Benes V., Garson J. a, Hellemans J., Huggett J., Kubista M., … Wittwer C.T. (2009) The MIQE guidelines: minimum information for publication of quantitative real-time PCR experiments. Clinical chemistry 55, 611–22, 10.1373/clinchem.2008.112797.

Casper B.B. & Jackson R.B. (1997) Plant Competition Underground. Annual Review of Ecology and Systematics 28, 545–570, 10.1146/annurev.ecolsys.28.1.545.

Chaumont F. & Tyerman S.D. (2014) Aquaporins: Highly Regulated Channels Controlling Plant Water Relations. Plant Physiology 164, 1600–1618, 10.1104/pp.113.233791.

Chen J., Tang H.-H., Li L., Qin S.-J., Wang G.-P., & Hong N. (2017) Effects of virus infection on plant growth, root development and phytohormone levels in in vitro-cultured pear plants. Plant Cell, Tissue and Organ Culture (PCTOC) 131, 359–368, 10.1007/s11240-017-1289-1.

Corrêa R.L., Sanz-Carbonell A., Kogej Z., Müller S.Y., Ambrós S., López-Gomollón S., … Elena S.F. (2020) Viral Fitness Determines the Magnitude of Transcriptomic and Epigenomic Reprograming of Defense Responses in Plants. Molecular Biology and Evolution 37, 1866–1881, 10.1093/molbev/msaa091.

Culver J.N. & Padmanabhan M.S. (2007) Virus-Induced Disease: Altering Host Physiology One Interaction at a Time. Annual Review of Phytopathology 45, 221–243, 10.1146/annurev.phyto.45.062806.094422.

De Vylder J., Vandenbussche F., Hu Y., Philips W., & Van Der Straeten D. (2012) Rosette Tracker: An Open Source Image Analysis Tool for Automatic Quantification of Genotype Effects. Plant Physiology 160, 1149–1159, 10.1104/pp.112.202762.

El Aou-ouad H., Pou A., Tomás M., Montero R., Ribas-Carbo M., Medrano H., & Bota J. (2017) Combined effect of virus infection and water stress on water flow and water economy in grapevines. Physiologia Plantarum 160, 171–184, 10.1111/ppl.12541.

Erion G.G. & Riedell W.E. (2012) Barley Yellow Dwarf Virus Effects on Cereal Plant Growth and Transpiration. Crop Science 52, 2794–2799, 10.2135/cropsci2012.02.0138.

Ertunç F. (2020) Physiology of virus-infected plants. In Applied Plant Virology. pp. 199–205. Elsevier.

Gibbs A.J., Hajizadeh M., Ohshima K., & Jones R.A.C. (2020) The Potyviruses: An Evolutionary Synthesis Is Emerging. Viruses 12, 132, 10.3390/v12020132.

Guo D.-P., Guo Y.-P., Zhao J.-P., Liu H., Peng Y., Wang Q.-M., … Rao G.-Z. (2005) Photosynthetic rate and chlorophyll fluorescence in leaves of stem mustard (Brassica juncea var. tsatsai) after turnip mosaic virus infection. Plant Science 168, 57–63, 10.1016/j.plantsci.2004.07.019.

Hall A.E. & Loomis R.S. (1972) An Explanation for the Difference in Photosynthetic Capabilities of Healthy and Beet Yellows Virus-infected Sugar Beets (*Beta vulgaris* L.). Plant Physiology 50, 576–580, 10.1104/pp.50.5.576.

Inoue-Nagata A.K., Jordan R., Kreuze J., Li F., López-Moya J.J., Mäkinen K., … ICTV Report Consortium (2022) ICTV Virus Taxonomy Profile: Potyviridae 2022: This article is part of the ICTV Virus Taxonomy Profiles collection. Journal of General Virology 103, 10.1099/jgv.0.001738.

Jež-Krebelj A., Rupnik-Cigoj M., Stele M., Chersicola M., Pompe-Novak M., & Sivilotti P. (2022) The Physiological Impact of GFLV Virus Infection on Grapevine Water Status: First Observations. Plants 11, 161, 10.3390/plants11020161.

Jones R.A.C. & Naidu R.A. (2019) Global Dimensions of Plant Virus Diseases: Current Status and Future Perspectives. Annual Review of Virology 6, 387–409, 10.1146/annurev-virology-092818-015606.

Lakens D. (2013) Calculating and reporting effect sizes to facilitate cumulative science: a practical primer for t-tests and ANOVAs. Frontiers in Psychology 4, 10.3389/fpsyg.2013.00863.

Laur J. & Hacke U.G. (2013) Transpirational demand affects aquaporin expression in poplar roots. Journal of Experimental Botany 64, 2283–2293, 10.1093/jxb/ert096.

Lindsey D.W. (1975) Effects of Maize Dwarf Mosaic Virus on Water Relations of Corn. Phytopathology 65, 434, 10.1094/Phyto-65-434.

Lo Gullo M.A., Nardini A., Salleo S., & Tyree M.T. (1998) Changes in root hydraulic conductance (*K*_R_) of *Olea oleaster* seedlings following drought stress and irrigation. New Phytologist 140, 25–31, 10.1046/j.1469-8137.1998.00258.x.

Loebenstein G. (2008) Plant Virus Diseases: Economic Aspects. In Encyclopedia of Virology. pp. 197–201. Elsevier.

López-González S., Gómez-Mena C., Sánchez F., Schuetz M., Samuels A.L., & Ponz F. (2021) The Effects of Turnip Mosaic Virus Infections on the Deposition of Secondary Cell Walls and Developmental Defects in Arabidopsis Plants Are Virus-Strain Specific. Frontiers in Plant Science 12, 741050, 10.3389/fpls.2021.741050.

Lunello P., Mansilla C., Sánchez F., & Ponz F. (2007) A Developmentally Linked, Dramatic, and Transient Loss of Virus from Roots of Arabidopsis thaliana Plants Infected by Either of Two RNA Viruses. Molecular Plant-Microbe Interactions® 20, 1589–1595, 10.1094/MPMI-20-12-1589.

Mäkinen K. (2023) How do they do it? The infection biology of potyviruses. In Advances in Virus Research. pp. 1–79. Elsevier.

Manacorda C.A. & Asurmendi S. (2018) Arabidopsis phenotyping through geometric morphometrics. GigaScience 7, giy073, 10.1093/gigascience/giy073.

Manacorda C.A., Gudesblat G., Sutka M., Alemano S., Peluso F., Oricchio P., … Asurmendi S. (2021) TUMV triggers stomatal closure but reduces drought tolerance in Arabidopsis. Plant, Cell & Environment 44, 1399–1416, 10.1111/pce.14024.

Manacorda C.A., Mansilla C., Debat H.J., Zavallo D., Sánchez F., Ponz F., & Asurmendi S. (2013) Salicylic Acid Determines Differential Senescence Produced by Two *Turnip mosaic virus* Strains Involving Reactive Oxygen Species and Early Transcriptomic Changes. Molecular Plant-Microbe Interactions® 26, 1486–1498, 10.1094/MPMI-07-13-0190-R.

Maurel C., Simonneau T., & Sutka M. (2010) The significance of roots as hydraulic rheostats. Journal of Experimental Botany 61, 3191–3198, 10.1093/jxb/erq150.

Maurel C., Verdoucq L., & Rodrigues O. (2016) Aquaporins and plant transpiration. Plant, Cell & Environment 39, 2580–2587, 10.1111/pce.12814.

Mirzayeva S., Huseynova I., Özmen C.Y., & Ergül A. (2023) Physiology and Gene Expression Analysis of Tomato (Solanum lycopersicum L.) Exposed to Combined-Virus and Drought Stresses. The Plant Pathology Journal 39, 466–485, 10.5423/PPJ.OA.07.2023.0103.

Mishra R., Shteinberg M., Shkolnik D., Anfoka G., Czosnek H., & Gorovits R. (2022) Interplay between abiotic (drought) and biotic (virus) stresses in tomato plants. Molecular Plant Pathology 23, 475–488, 10.1111/mpp.13172.

Moshelion M., Halperin O., Wallach R., Oren R., & Way D.A. (2015) Role of aquaporins in determining transpiration and photosynthesis in water-stressed plants: crop water-use efficiency, growth and yield. Plant, Cell & Environment 38, 1785–1793, 10.1111/pce.12410.

Murray R.R., Emblow M.S.M., Hetherington A.M., & Foster G.D. (2016) Plant virus infections control stomatal development. Scientific Reports 6, 34507, 10.1038/srep34507.

Nellist C.F., Ohshima K., Ponz F., & Walsh J.A. (2022) Turnip mosaic virus, a virus for all seasons. Annals of Applied Biology 180, 312–327, 10.1111/aab.12755.

Ozu M., Galizia L., Acuña C., & Amodeo G. (2018) Aquaporins: More Than Functional Monomers in a Tetrameric Arrangement. Cells 7, 209, 10.3390/cells7110209.

Passioura J.B. (1982) Water in the Soil-Plant-Atmosphere Continuum. In Physiological Plant Ecology II. (eds O.L. Lange, P.S. Nobel, C.B. Osmond, & H. Ziegler), pp. 5–33. Springer Berlin Heidelberg, Berlin, Heidelberg.

Peng Y., Dallas M.M., Ascencio-Ibáñez J.T., Hoyer J.S., Legg J., Hanley-Bowdoin L., … Yin H. (2022) Early detection of plant virus infection using multispectral imaging and spatial–spectral machine learning. Scientific Reports 12, 3113, 10.1038/s41598-022-06372-8.

Pfaffl M.W., Horgan G.W., & Dempfle L. (2002) Relative expression software tool (REST©) for group-wise comparison and statistical analysis of relative expression results in real-time PCR.

Pinheiro & Bates (2000) Mixed-Effects Models in S and S-PLUS. Springer-Verlag, New York.

Pollari M., Sipari N., Poque S., Himanen K., & Mäkinen K. (2022) Effects of Poty-Potexvirus Synergism on Growth, Photosynthesis and Metabolite Status of *Nicotiana benthamiana*. Viruses 15, 121, 10.3390/v15010121.

Prasch C.M. & Sonnewald U. (2013) Simultaneous Application of Heat, Drought, and Virus to Arabidopsis Plants Reveals Significant Shifts in Signaling Networks. Plant Physiology 162, 1849–1866, 10.1104/pp.113.221044.

Qin L., Ding S., & He Z. (2023) Compositional biases and evolution of the largest plant RNA virus order Patatavirales. International Journal of Biological Macromolecules 240, 124403, 10.1016/j.ijbiomac.2023.124403.

R Core Team (2020) R: A language and environment for statistical computing.

Ramakers C., Ruijter J.M., Deprez R.H.L., & Moorman A.F.M. (2003a) Assumption-free analysis of quantitative real-time polymerase chain reaction (PCR) data. Neuroscience Letters 339, 62–66, 10.1016/S0304-3940(02)01423-4.

Ramakers C., Ruijter J.M., Lekanne Deprez R.H., & Moorman A.F.M. (2003b) Assumption-free analysis of quantitative real-time polymerase chain reaction (PCR) data. Neuroscience Letters 339, 62–66, 10.1016/S0304-3940(02)01423-4.

Revers F. & García J.A. (2015) Molecular Biology of Potyviruses. In Advances in Virus Research. pp. 101–199. Elsevier.

Di Rienzo J.A. (2009) fgStatistics. Statistical software for the analysis of microarray data.

Rodríguez M., Muñoz N., Lenardon S., & Lascano R. (2012) The chlorotic symptom induced by Sunflower chlorotic mottle virus is associated with changes in redox-related gene expression and metabolites. Plant Science 196, 107–116, 10.1016/j.plantsci.2012.08.008.

Rodriguez-Dominguez C.M. & Brodribb T.J. (2020) Declining root water transport drives stomatal closure in olive under moderate water stress. New Phytologist 225, 126–134, 10.1111/nph.16177.

RStudio Team (2019) RStudio: Integrated Development for R.

Sade N., Galkin E., & Moshelion M. (2015) Measuring Arabidopsis, Tomato and Barley Leaf Relative Water Content (RWC). BIO-PROTOCOL 5, 10.21769/BioProtoc.1451.

Sade N., Gebretsadik M., Seligmann R., Schwartz A., Wallach R., & Moshelion M. (2009) The Role of Tobacco Aquaporin1 in Improving Water Use Efficiency, Hydraulic Conductivity, and Yield Production Under Salt Stress. Plant Physiology 152, 245–254, 10.1104/pp.109.145854.

Sánchez F., Manrique P., Mansilla C., Lunello P., Wang X., Rodrigo G., … Ponz F. (2015) Viral Strain-Specific Differential Alterations in *Arabidopsis* Developmental Patterns. Molecular Plant-Microbe Interactions® 28, 1304–1315, 10.1094/MPMI-05-15-0111-R.

Sánchez F., Martínez-Herrera D., Aguilar I., & Ponz F. (1998) Infectivity of turnip mosaic potyvirus cDNA clones and transcripts on the systemic host Arabidopsis thaliana and local lesion hosts. Virus Research 55, 207–219, 10.1016/S0168-1702(98)00049-5.

Schenk H.J. & Jackson R.B. (2002) Rooting depths, lateral root spreads and below-ground/above-ground allometries of plants in water-limited ecosystems. Journal of Ecology 90, 480–494, 10.1046/j.1365-2745.2002.00682.x.

Scholthof K.G., Adkins S., Czosnek H., Palukaitis P., Jacquot E., Hohn T., … Foster G.D. (2011) Top 10 plant viruses in molecular plant pathology. Molecular Plant Pathology 12, 938–954, 10.1111/j.1364-3703.2011.00752.x.

Steudle E. (2000) Water uptake by roots: effects of water deficit. Journal of Experimental Botany 51, 1531–1542, 10.1093/jexbot/51.350.1531.

Sutka M., Li G., Boudet J., Boursiac Y., Doumas P., & Maurel C. (2011a) Natural Variation of Root Hydraulics in Arabidopsis Grown in Normal and Salt-Stressed Conditions. Plant Physiology 155, 1264–1276, 10.1104/pp.110.163113.

Sutka M., Li G., Boudet J., Boursiac Y., Doumas P., & Maurel C. (2011b) Natural Variation of Root Hydraulics in Arabidopsis Grown in Normal and Salt-Stressed Conditions. Plant Physiology 155, 1264–1276, 10.1104/pp.110.163113.

Sutka M.R., Manzur M.E., Vitali V.A., Micheletto S., & Amodeo G. (2016) Evidence for the involvement of hydraulic root or shoot adjustments as mechanisms underlying water deficit tolerance in two Sorghum bicolor genotypes. Journal of Plant Physiology 192, 13–20, 10.1016/j.jplph.2016.01.002.

Torres-Ruiz J.M., Cochard H., Delzon S., Boivin T., Burlett R., Cailleret M., … Martin-StPaul N.K. (2024) Plant hydraulics at the heart of plant, crops and ecosystem functions in the face of climate change. New Phytologist 241, 984–999, 10.1111/nph.19463.

Tournaire-Roux C., Sutka M., Javot H., Gout E., Gerbeau P., Luu D.-T., … Maurel C. (2003) Cytosolic pH regulates root water transport during anoxic stress through gating of aquaporins. Nature 425, 393–397, 10.1038/nature01853.

Turner N.C. (1981) Techniques and experimental approaches for the measurement of plant water status. Plant and Soil 58, 339–366, 10.1007/BF02180062.

Vadez V. (2014) Root hydraulics: The forgotten side of roots in drought adaptation. Field Crops Research 165, 15–24, 10.1016/j.fcr.2014.03.017.

Vaisman M., Hak H., Arazi T., & Spiegelman Z. (2023) The Impact of Tobamovirus Infection on Root Development Involves Induction of *Auxin Response Factor 10a* in Tomato. Plant and Cell Physiology 63, 1980–1993, 10.1093/pcp/pcab179.

Vaissie P., Monge A., & Husson F. (2015) Factoshiny: Perform Factorial Analysis from “FactoMineR” with a Shiny Application. 2.7, 10.32614/CRAN.package.Factoshiny.

Vandeleur R.K., Sullivan W., Athman A., Jordans C., Gilliham M., Kaiser B.N., & Tyerman S.D. (2014) Rapid shoot-to-root signalling regulates root hydraulic conductance via aquaporins. Plant, Cell & Environment 37, 520–538, 10.1111/pce.12175.

Villordon A.Q. & Clark C.A. (2014) Variation in Virus Symptom Development and Root Architecture Attributes at the Onset of Storage Root Initiation in ‘Beauregard’ Sweetpotato Plants Grown with or without Nitrogen. PLoS ONE 9, e107384, 10.1371/journal.pone.0107384.

Vitali V., Bellati jorge, & Soto G. (2015) Root hydraulic conductivity and adjustments in stomatal conductance: hydraulic strategy in response to salt stress in a halotolerant species.

Vitti A., Nuzzaci M., Scopa A., Tataranni G., Remans T., Vangronsveld J., & Sofo A. (2013) Auxin and Cytokinin Metabolism and Root Morphological Modifications in Arabidopsis thaliana Seedlings Infected with Cucumber mosaic virus (CMV) or Exposed to Cadmium. International Journal of Molecular Sciences 14, 6889–6902, 10.3390/ijms14046889.

Walsh J.A. & Jenner C.E. (2002) *Turnip mosaic virus* and the quest for durable resistance. Molecular Plant Pathology 3, 289–300, 10.1046/j.1364-3703.2002.00132.x.

Wang A. (2021) Cell-to-cell movement of plant viruses via plasmodesmata: a current perspective on potyviruses. Current Opinion in Virology 48, 10–16, 10.1016/j.coviro.2021.03.002.

Xu P., Chen F., Mannas J.P., Feldman T., Sumner L.W., & Roossinck M.J. (2008) Virus infection improves drought tolerance. New Phytologist 180, 911–921, 10.1111/j.1469-8137.2008.02627.x.

Yang C., Guo R., Jie F., Nettleton D., Peng J., Carr T., … Whitham S.A. (2007) Spatial Analysis of *Arabidopsis thaliana* Gene Expression in Response to *Turnip mosaic virus* Infection. Molecular Plant-Microbe Interactions® 20, 358–370, 10.1094/MPMI-20-4-0358.

Ye J., Coulouris G., Zaretskaya I., Cutcutache I., Rozen S., & Madden T.L. (2012) Primer-BLAST: A tool to design target-specific primers for polymerase chain reaction. BMC Bioinformatics 13, 134, 10.1186/1471-2105-13-134.

